# p53-induced GDF-15 expression promotes a pro-regenerative response in human chondrocytes upon cartilage injury

**DOI:** 10.1101/2024.04.10.588843

**Authors:** Sara Sofi Marques, Alexandra Liebaug, Svenja Maurer, Dietrich Rothenbacher, Rolf E Brenner, Jana Riegger

## Abstract

Posttraumatic osteoarthritis (PTOA) is a special form of OA, developing after joint injuries. Except for some minor differences between the clinical characteristics of patients suffering from idiopathic OA (IOA) and PTOA, no biologic marker has yet been identified to distinguish IOA from PTOA. In this study, we investigated the expression of the stress-responsive cytokine growth differentiation factor 15 (GDF-15) in clinical samples from the Ulm OA study cohort (PTOA: n =12; IOA: n=54) and in a human ex vivo cartilage trauma model.

We found significantly higher GDF-15 levels in synovial fluid of end-stage PTOA patients as compared to IOA patients. Further, we confirmed that fibroblast-like synoviocytes secreted increased levels of GDF-15 after stimulation with medium of *ex vivo*-traumatized cartilage. GDF-15 and its receptor, GFRAL, were significantly elevated in highly degenerated OA cartilage. By means of a human cartilage trauma model, we discovered that chondrocytes produced GDF-15 upon tissue injury, while antioxidative treatment attenuated GDF-15 secretion. We confirmed that oxidative stress and subsequent activation of p53 resulted in GDF-15 expression. As a transcriptional target of p53, GDF-15 was associated with chondrosenescence. Although the cytokine might therefore represent a senescence-associated secretory phenotype (SASP) factor, stimulation with exogenous GDF-15 did not cause detrimental effects but induced a pro-regenerative response in chondrocytes, characterized by enhanced proliferation as well as chondro- and cell protection after cartilage trauma. Overall, this study first describes GDF-15 as a stress-responsive and potentially pro-regenerative cytokine in the context of human PTOA.

## Introduction

Osteoarthritis (OA) is commonly considered as the most abundant joint disease and major cause of disability in elderly people. Most cases of OA develop without a certain cause, but seem to be associated with gender, age, and genetic predisposition. This form of OA is also referred to as primary or idiopathic OA (IOA). In contrast to IOA, posttraumatic OA (PTOA) results from a certain event, namely a preceding joint injury, such as meniscal or ligament rupture, chondral defects, and articular fracture [1]. Although there are some epidemiological differences between IOA and PTOA, for example the earlier onset of PTOA, it remains largely unclear whether it is possible to distinguish the pathogenesis of the different forms on molecular or biochemical level [2]. PTOA accounts for about 12 % of all OA cases and represents a particularly severe burden due to the significantly younger age of the affected patients, which is accompanied by early invalidity and thus inability to work, as well as enhanced lifetime risk for revision arthroplasty [3]. Not all patients, receiving a joint injury, are deemed to develop a PTOA in later life, because further factors are thought to be involved in disease progression [4]. Identification of novel biomarkers serving as prognostic tools to estimate the actual risk of PTOA development after injury or as therapeutic target structures, is of high priority.

Recently, we compared the clinical parameters of patients suffering from knee IOA or PTOA among the Ulm Osteoarthritis Study cohort. The study revealed differences between IOA and PTOA patients regarding the WOMAC pain subscale and general physical function at baseline (before knee joint replacement), as well as serum high-sensitivity cardiac troponin T levels and mortality (manuscript in preparation). In the present study, we focused on growth differentiation factor 15 (GDF-15) as a potential indicator to differentiate IOA and PTOA. The protein has previously been described as elevated in different forms of rheumatic diseases, including systemic lupus erythematosus, spondyloarthritis, and rheumatoid arthritis, in which it was associated with disease severity and activity [5-7]. GDF-15, also referred to as the macrophage inhibitory cytokine 1, is a stress-inducible cytokine and atypical member of the transforming growth factor-β (TGF-β) superfamily, which has emerged as a marker of all-cause mortality in heart failure and cancer, among others [8, 9]. Accordingly, we previously reported that the cytokine represents a strong and independent risk factor for decreased survival in subjects undergoing unilateral total hip or knee arthroplasty due to advanced OA within the Ulm OA study cohort [10]. Despite its bad reputation as all-cause mortality marker, increased GDF-15 expression has also been discussed as a survival response [11]. Accordingly, GDF-15 is produced by various cell types during acute tissue injury and was described as prominent senescence-associated secretory phenotype (SASP) factor in senescent cells [12]. Although transient senescence is considered as beneficial in terms of tissue remodelling and repair, chronic accumulation of senescent cells in cartilage or other tissues contributes to progressive degeneration and dysfunction due to the excessive secretion of catabolic and pro-inflammatory SASP factors [13, 14]. Considering its association with injury response and senescence, mechanisms which are obviously involved in the pathogenesis of PTOA [15], we hypothesized that GDF-15 might be a potential mediator in trauma-induced cartilage degeneration.

In the present study, we first confirmed significantly higher concentrations of GDF-15 in clinical synovial fluid samples of PTOA patients as compared to IOA patients. Moreover, we found clear evidence that GDF-15 was expressed by chondrocytes in highly degenerated human cartilage as well as after *ex vivo* trauma. Further investigation revealed that GDF-15 in human articular chondrocytes (hAC) was promoted by oxidative stress and subsequent activation of p53. Strikingly, we observed that GDF-15 secretion did not induce pathophysiologic processes but contributed to cell proliferation and protection, implying potentially pro-regenerative effects after cartilage injury. Taken together, this is the first study on GDF-15 in PTOA considering mainly clinical samples and providing a novel insight into the role of this stress-inducible cytokine.

## Material and Methods

### Clinical samples from the Ulm Osteoarthritis Study

Serum, synovial fluid, and synovial tissue samples were collected in the course of the longitudinal Ulm Osteoarthritis Study from patients suffering from advanced OA of the knee as determined by radiographic changes of grade ≥2 according to Kellgren and Lawrence [16], and undergoing unilateral total knee joint replacement (TKR) between January 1995 and December 1996 and stored at −80°C until use. The inclusion criteria (i.e., Caucasian, age <76 years, absence of malignancies, inflammatory diseases, rheumatoid arthritis, or corticosteroid medication; no previous joint replacement) were fulfilled by 54 patients. The patients were assigned in two groups: IOA patients (n= 42) and PTOA (n= 12), which included traumatic meniscus lesion/ ligament rupture/ knee joint luxation, intraarticular fractures (knee joint, patella, tibia head), general severe knee injury (with joint effusion, hemarthros etc.). Relevant clinical characteristics of patients included in this study are described in **Table 1**. Serum concentrations of cystatin C, COMP, and hs-CRP were detected as previously described [10, 17]. The study was approved by the local Ethics Committee of the University Ulm (No. 40/94 and 164/14) and written informed consent of the patients was given.

**Table 1:** Clinical characteristics of patients with PTOA and IOA at the time point of knee joint replacement.

| Parameter | Category/<br>Statistics | Idiopathic OA<br>(N = 42) | Posttraumatic OA<br>(N= 12) | Total<br>(N= 54) | P Value |
| --- | --- | --- | --- | --- | --- |
| Age (years) | Mean (SD) | Mean: 66.3 | Mean: 66.5 | Mean: 66.4 | 0.873 |
| Sex | Male<br>Female | Male: 13 (30.9 %)<br>Female: 29 (69.0 %) | Male: 2 (16.6 %)<br>Female: 10 (83.3 %) | Male: 15 (27.7 %)<br>Female: 39 (72.2 %) | 0.260 |
| BMI (kg/m<sup>2</sup>) | Mean (SD) | Mean: 29.8 | Mean: 28.9 | Mean: 29.6 | 0.603 |
| Diabetes<br>Mellitus | Yes<br>No | 6 (13.9 %)<br>36 (85.7 %) | 2 (16.6 %)<br>10 (83.3 %) | 8 (14.8 %)<br>40 (85.1 %) | 0.695 |

### Chondrocyte culture and *ex vivo* cartilage trauma model

Human cartilage was obtained from OA patients undergoing TRK with informed consent according to the regulations of the ethics committee of the University of Ulm (164/14). For the *ex vivo* cartilage trauma model and chondrocyte isolation, only macroscopically intact cartilage (OARSI ≤ 1) was used.

Human chondrocytes were isolated by enzymatic digestion. Cartilage was minced and pre-digested with 0.2% pronase for 45 minutes, followed by an overnight incubation with 0.025% collagenase at 37°C in a rotator. Chondrocytes were cultured in chondrocyte medium (CM) containing 10% FCS and used at passages ≤ 2. Experiments were performed under serum-reduced conditions (1:1 CM and serum-free medium (SFM)).

*Ex vivo* traumatization of cartilage explants was performed as previously described. In brief, 6 mm diameter full-thickness cartilage explants were prepared. After a resting time of 24 h, explants were impacted by means of our drop tower device with an energy of 0.59 J [18-20]. Tissue experiments were performed under serum-free conditions in SFM. Cartilage tissue and chondrocytes were cultured at 37°C, 5% CO2, and 95% humidity.

### Human FLS experiments

Human fibroblast-like synoviocytes (FLS) were isolated as previously described [20]. In short, synovial tissue of 6 patients was minced and incubated in 25 mg collagenase type VIII for 90 min at 37°C, filtered (70 μm), and cultured in FLS serum-containing medium. Cells were used at passages 2-4. After adherence, cells were stimulated with conditioned medium from unimpacted or impacted cartilage (see above), diluted 1:1 in SFM, and cultured at 27 °C or 37 °C for 4 days.

### Medium composition

Chondrocyte medium (CM): 1:1 DMEM (1 g/L glucose)/ Ham’s F12, 10% heat-inactivated fetal calf serum (FCS), 0.5% penicillin/streptomycin, 2 mM L-glutamine, 10 μg/mL 2-phospho-L-ascorbic acid trisodium salt.

Serum-free medium (SFM): DMEM (1 g/L glucose), 1% sodium pyruvate, 0.5% L-glutamine, 1% non-essential amino acids, 0.5% penicillin/streptomycin, 10 μg/ml 2-phospho-L-ascorbic acid trisodium salt, 0.1% insulin-transferrin-selenium (ITS).

FLS serum-containing medium: High glucose (4.5 g/L) DMEM, 10% fetal bovine serum, 0.5% L-glutamine, 0.5% penicillin/streptomycin (Sigma-Aldrich).

### Gene expression analysis

In case of cartilage samples, tissue was snap frozen in liquid nitrogen and pulverized with a microdismembrator S (B. Braun Biotech, Melsungen, Germany) before total RNA isolation by means of the Lipid Tissue Mini Kit (Qiagen, Hilden, Germany). Isolated cells were lysed in RLT buffer, followed by total mRNA isolation using the Qiagen Mini Kit. RNA was reverse transcribed with the Omniscript RT Kit (Qiagen), according to the manufacturer`s protocol.

Quantitative real-time PCR analysis was performed by using a StepOnePlus™ Real-Time PCR System (Applied Biosystems, Darmstadt, Germany). Target transcripts were analysed by TaqMan® Gene Expression Assays or self-designed primers and the respective Master Mix, as indicated in **Table S1**.

In brief, relative expression levels of target genes were determined using the 2^−ΔΔCt^ method, by normalization to the endogenous controls (18S rRNA, GAPDH and HPRT1) as previously described [18].

### alamarBlue Assay

The alamarBlue assay (BioRad, Munich, Germany) was used to measure the metabolic activity of living cells in order to determine proliferation/ cytotoxicity. In brief, cells were incubated for 3 h in a 5% alamarBlue solution in serum-reduced medium at 37 °C. The emitted fluorescence was detected at a 550 nm excitation and 590 nm emission by using the multimode microplate reader Infinite M200 Pro (Tecan Deutschland, Crailsheim, Germany).

### Scratch Assay

A scratch assay, also known as wound healing or migration assay, was used to assess migratory activity of chondrocytes in presence of GDF-15. A straight scratch was applied in a confluent chondrocyte monolayer by using the tip of a 200 μL sterile pipette scraped across the surface. Images were captured after 0, 24, and 48 hours.

### GDF-15 ELISA

To quantify GDF-15 in synovial fluid samples and cell culture supernatants, the human GDF-15 ELISA Kit by R&D Systems was used, according to the manufacturer`s protocol. In brief, 100 µL of undiluted samples were applied. In case of synovial fluid, samples were subjected to a hyaluronidase digest (4 mg/mL; 1:1 mixed with sample) for 60 min at 37 °C. Influence of the hyaluronidase digest on GDF-15 detection is provided in **Figure S1A**.

### Immunohistochemistry

For IHC staining, cartilage and synovial tissue was fixed (4% paraformaldehyde) and embedded in paraffin. In brief, tissue sections were dewaxed and rehydrated, followed by H_2_O_2_ incubation for 30 min. For antigen retrieval, sections were incubated in citrate buffer (pH 6.0) at 60 °C overnight. Primary antibodies were used as follows: GDF-15 (Novus Biologicals, Wiesbaden, Germany; NBP1-81050; 1:100), GFRAL (Invitrogen, PA5-24545; 1:100), p53 (LS-Bio, Lynnwood, WA, US; LS-B7723; 1:100), p21 (Thermo Fisher Scientific, MA5-14949; 1:50), incubated overnight at 4 °C. The staining was performed with the Dako LSAB2 System-HRP kit (Dako, Glostrup, Denmark). Subsequently, cell nuclei were counter stained by Gill’s haematoxylin No. 3 (Sigma-Aldrich). Documentation was performed with an Axioskop 2 mot plus (Zeiss, Oberkochen, Germany).

### Immunofluorescence Staining of p53

hAC were fixed with formalin, permeabilized with 0.1% Triton, and incubated for 1 h at 37 °C with blocking buffer (Agilent Technologies, Waldbronn, Germany). Afterwards, cells were stained with anti-p53 (LSBio, LS-B7723; 1:250) for 3 h at RT, followed by an incubation with a secondary antibody (abcam, ab150077, Alexa 488, 1:500) for 30 min at RT. Nuclei were counterstained with 0.25 µg/mL Dapi for 15 min. Area, fluorescence, and integrated density of the cells were measured using Fiji (Version 2.1.0/1.53c; open-source software). To distinguish between specific and nonspecific signals, the average fluorescence of untreated hACs was calculated and then subtracted from the measured fluorescence of each cell. Subsequently, the corrected total cell fluorescence (CTCF) was determined for the p53-positive cells.

### DCFDA Assay

Analysis of cytoplasmatic ROS levels was performed by means of the DCFDA/H2DCFDA-Cellular ROS Assay Kit (Abcam). In short, cultured hAC were incubated with a 1 µM DCFDA working solution for 45 min at 37 °C. Afterward, samples were analyzed with a fluorescence microscope. The CTCF was assessed as described above.

### Live/dead cell cytotoxicity assay

To determine the percentage of viable cells, a Live/Dead® Viability/Cytotoxicity Assay (Molecular Probes, Invitrogen) was performed. Unfixed tissue sections (0.5 mm thickness) were stained with 1 μM calcein AM and 2 μM ethidium homodimer-1 for 30 min. After washing in PBS, they were microscopically analysed by means of a z-stack module (software AxioVision, Carl Zeiss, Jena, Germany).

### DMMB and Griess Assay

Content of proteoglycan release into culture media (μg/mL) was quantified using the photometric 1,9-dimethylmethylene blue assay as previously described [18]. Analysis is based upon the binding of sulfated glycosaminoglycan (GAG) in the presence of 0.24 M GuHCl. In case of NO release, levels were determined by quantification of nitrite, a stable end product of the NO metabolism, using a Griess assay (Griess Reagent System; Promega) [21].

### RNA sequencing

Total RNA from cells was extracted as described above. Quality assessment, library preparation, RNA-sequencing, and bioinformatic analysis was carried out by Biomarker Technologies (BMK; Münster, Germany). Libraries were sequenced using an Illumina Novaseq 6000 (PE150) system.

### Statistical Analysis

Data were analysed using GraphPad Prism9 (GraphPad Software, Inc., San Diego, CA, USA). Datasets with n ≥ 5 were tested for outliers by means of the Grubbs’ outlier test. Outliers were not included in the statistical analyses. Each data point represents an independent biological replicate (donor). Data sets of two groups were analysed by means of a two-tailed (multiple) t-test. In case of data sets with three groups and more, a one-way ANOVA with Sidak post-hoc test (parametric distribution) or Kruskal-Wallis with a Dunn’s post-hoc test (non-parametric distribution) was used. Association between two parameters was analysed by means of a Pearson correlation analysis. In each case, significance level was set to α = 0.05.

## Results

### Synovial GDF-15 concentrations are significantly higher in PTOA as compared to IOA samples

To prove our hypothesis that GDF-15 is differentially expressed in IOA and PTOA patients, we analyzed clinical serum and synovial fluid samples derived from the Ulm Osteoarthritis study cohort. Although we found increased serum levels of GDF-15 (sGDF-15) in the PTOA group, there was no significant difference compared to the IOA group (**Figure 1A**). To clarify whether GDF-15 was locally expressed in the affected knee joints of the respective patients, we investigated the synovial GDF-15 (syGDF-15) concentrations. In fact, syGDF-15 levels were significantly higher in synovial fluid samples of PTOA patients as compared to the IOA group (mean ± SD: PTOA 625.9 ng/mL ± 210.2; IOA 452.8 ng/mL ± 150.3; **Figure 1B**). While syGDF-15 concentrations significantly correlated with the serum GDF-15 levels in the overall collective, we could not find any association in the IAO group and only a trend among the PTOA group (**Figure 1C**). Moreover, syGDF-15 levels were significantly associated with serum cystatin C and serum COMP in the overall cohort (**Figure 1D**). Further, syGDF-15 was associated with serum cystatin C in both, the IOA and PTOA group, whereas significant correlations between syGDF-15 and age or hs-CRP were only found in PTOA patients (**Figure 1D**). No significant differences in syGDF-15 levels were observed between female and male patients (**Supplement S1B**).

**Figure 1:**
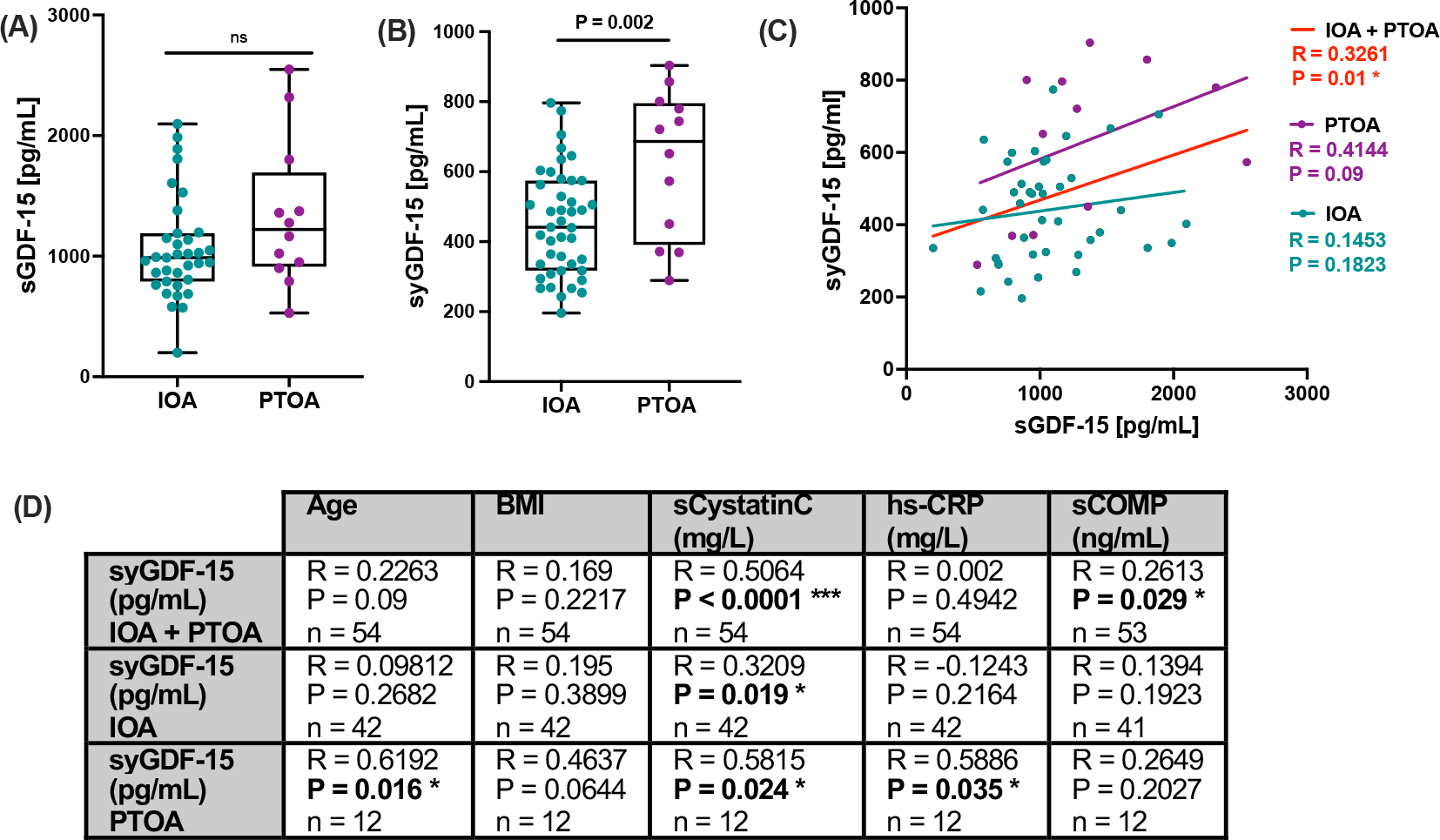
Assessment of local GDF-15 levels in synovial fluid of PTOA and IOA patients. (A) Serum GDF-15 (sGDF-15) and (B) synovial GDF-15 (syGDF-15) concentrations of IOA and PTOA patients at the time point of knee TEP implantation were quantified by means of ELISAs. (C) Corresponding correlation analysis between syGDF-15 and sGDF-15 in IOA and PTOA patients. (C) Overview of correlation analyses between syGDF-15 concentrations with clinical and laboratory parameters in IOA and PTOA patients separately or combined. sCOMP= serum COMP, sCystatinC= serum cystatin C.

Systemic hs-CRP levels were previously associated with synovial inflammation in OA patients [22], whereas serum cystatin C concentrations were found to correlate with disease activity and chronic inflammation in rheumatic arthritis [23]. Thus, our findings might indicate a potential connection between GDF-15 and local inflammatory processes in the PTOA-affected knee.

### Synovial cells express enhanced levels of GDF-15 in response to trauma-conditioned medium

To investigate the origin of syGDF-15 and its potential connection to synovitis, we quantified GDF-15 expressing cells in the synovial tissue of IOA and PTOA patients by immunostaining. As expected for IOA and PTOA, which are considered as low-grade inflammatory forms of arthritic disease [24], the histologic assessment revealed only mild synovitis and no difference between IOA and PTOA. Overall, GDF-15 expressing cells could be found in the synovial tissue of both groups and there was no clear association with the posttraumatic history of the patients or the syGDF-15 concentrations. Furthermore, GDF-15 was not co-located to infiltrated immune cells (**Figure 2A**).

**Figure 2:**
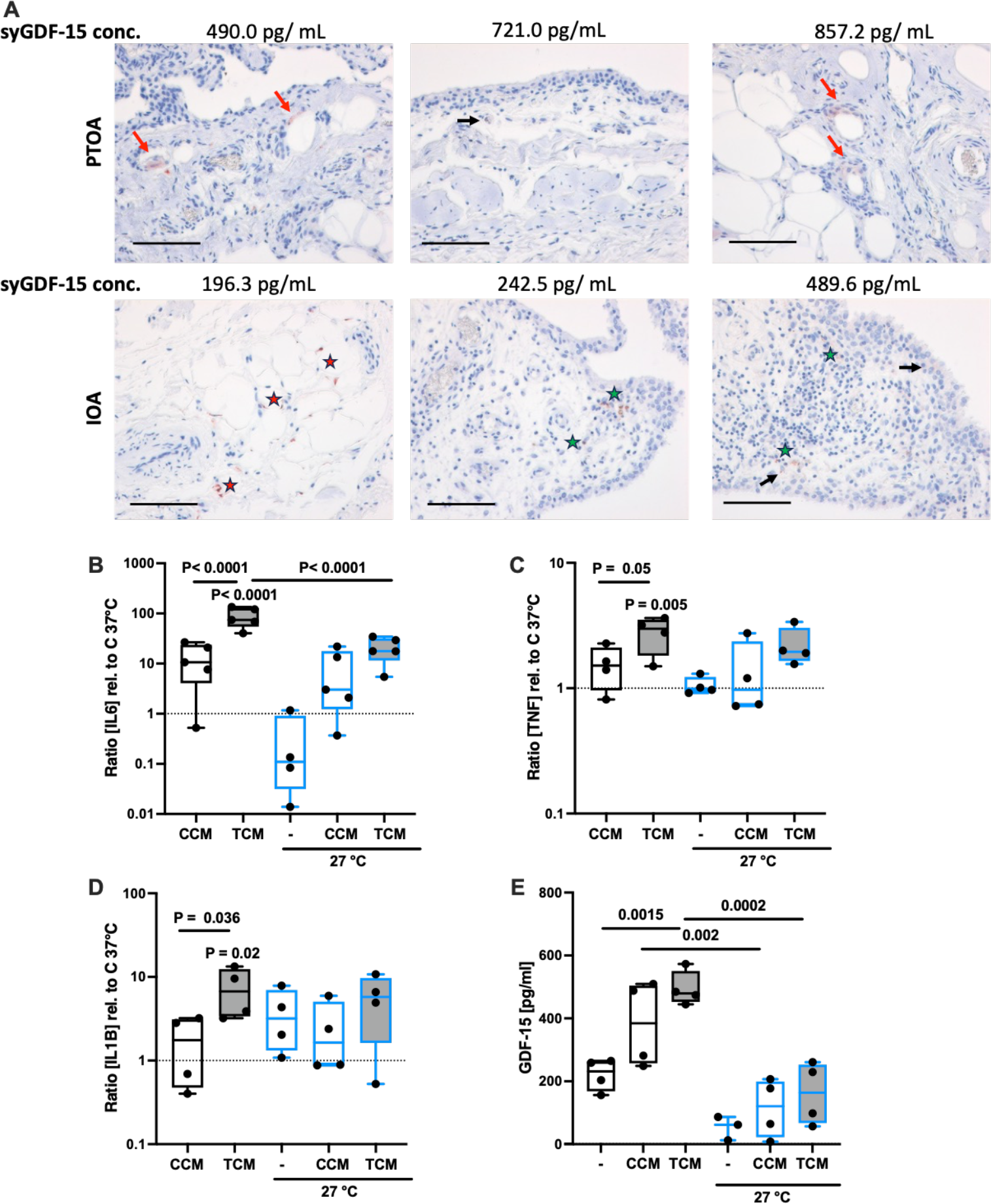
Expression of GDF-15 in synovial cells. (A) Representative images of IHC staining against GDF-15 in the synovial membrane of IOA and PTOA patients, including the respective syGDF-15 concentrations. GDF-15 expressing cells are highlighted as follows: synovial lining cells = black arrows; infiltrated immune/ small follicle-like lymphocytic cells = green star; endothelial cells = red arrow; adipocytes = red stars. The black bar represents 100 µm. (B) Gene expression analysis of IL6, (C) TNF, and (D) IL1B, as well as (E) release of GDF-15 by human fibroblast-like synoviocytes (hFLS) stimulated with cartilage-conditioned medium (CCM) or trauma-conditioned medium (TCM) and cultivated under normothermic (37 °C) or hypothermic (27 °C) conditions for 4d.

Previously, we reported a pro-inflammatory response of isolated hFLS after stimulation with trauma-conditioned medium (TCM) derived from *ex vivo* traumatized human cartilage explants. This activation is most likely triggered by the binding of trauma-derived damage-associated molecular patterns (DAMPs) to cellular pattern recognition receptors, in particularly toll-like receptors, and was found to be partly attenuated at hypothermic conditions (27 °C) [20]. We used this *in vitro* model to investigate whether activated hFLS express GDF-15. While the gene expression of IL6, TNF, and IL1B was significantly enhanced in hFLS upon TCM stimulation, only that of IL6 was significantly reduced at hypothermic conditions (**Figure 2B-D**). In line to the other cytokines, release of GDF-15 was significantly increased by hFLS in response to TCM. This effect was completely suppressed at 27 °C (**Figure 2E**).

Altogether, the results imply that hFLS might secrete GDF-15 in response to trauma-associated DAMPS.

### GDF-15 expression is significantly enhanced in chondrocytes of highly degenerated or traumatized cartilage

To clarify whether hAC contribute to syGDF-15 levels and might represent possible recipient cells of the cytokine, we investigated the expression of GDF-15 and its receptor GFRAL in macroscopically intact (OARSI ≤ 1) and highly degenerated (OARSI ≥ 3) cartilage of the same OA patients. While the gene expression of GDF15 was significantly increased in highly degenerated cartilage tissue, no difference was observed for GFRAL as compared to macroscopically intact tissue (**Figure 3A**). However, on protein level both GDF-15 and GFRAL were found to be significantly elevated in highly degenerated cartilage, as determined by IHC (**Figure 3B-D**). This finding indicates that GDF-15 expression is associated with cartilage degeneration and that osteoarthritic chondrocytes might be more susceptible to the cytokine due to the upregulation of GFRAL.

**Figure 3:**
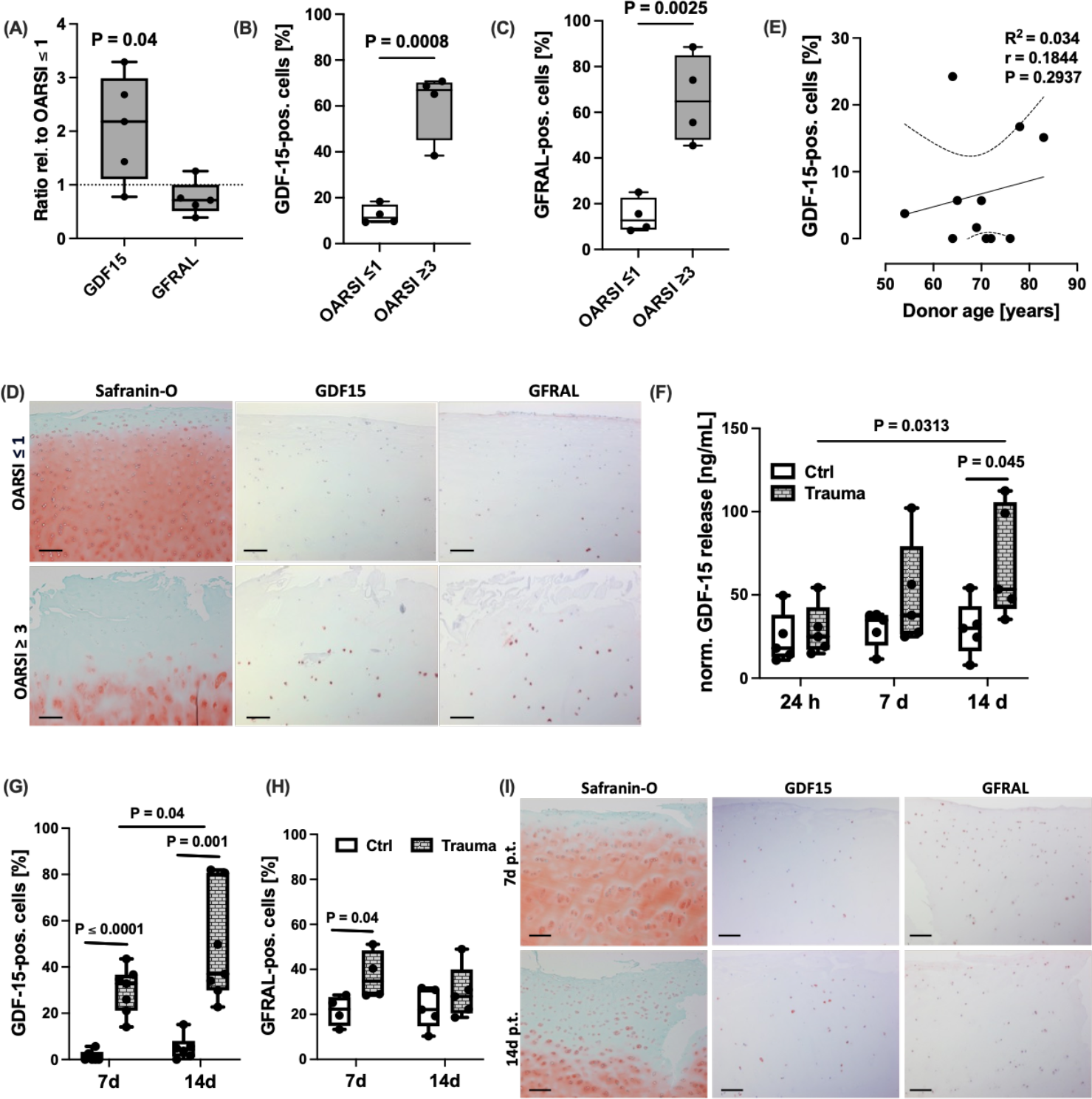
Expression of GDF-15 in highly degenerated and traumatized cartilage tissue. (A) Gene expression analysis of GDF15 and its receptor GFRAL in highly degenerated cartilage (OARSI ≥ 3) relative to macroscopically intact (OARSI ≤ 1) tissue. (B) Quantification of GDF-15 and (C) GFRAL expressing cells in macroscopically intact and highly degenerated cartilage by IHC. (D) Representative images of Safranin-O staining as well as IHC staining against GDF-15 and GFRAL in the respective tissue. (E) Correlation analysis between percentage of GDF-15 positive cells in macroscopically intact tissue and the respective donor age. (F) Release of GDF-15 at 24 h, 7 d, and 14 d after *ex vivo* cartilage trauma. Quantification of IHC staining against (G) GDF-15 and (H) GFRAL at 7 d and 14 d after ex vivo cartilage trauma. (I) Representative images of Safranin-O staining as well as IHC staining against GDF-15 and GFRAL in the respective tissue. The black bar in (D) and (I) represents 200 µm.

Although syGDF-15 concentration correlated with donor age of PTOA patients, as described above, we did not find an association between the percentage of GDF15-expressing chondrocytes in macroscopically intact cartilage tissue and age of the respective donors (**Figure 3E**).

Next, we wanted to clarify the potential link between cartilage injury and GDF-15 expression. For this purpose, human macroscopically intact cartilage was traumatized by means of an *ex vivo* drop tower device [18]. On gene expression level, GDF15 was induced 14 d after trauma, but not at the earlier time points (**Supplement S2A**), which was largely consistent with the time-dependent increase of GDF-15 biosynthesis as confirmed by ELISA and IHC (**Figure 3F,G,I**). GFRAL expression, in turn, was elevated 7 d post trauma, but was largely restored after 14 d (**Figure 3H,I**; **Supplement S2B**).

Overall, these results imply that GDF-15 expression is associated with cartilage injury and might occur as a delayed – not immediate – response.

### Expression of GDF-15 is regulated by stress-induced p53 activity

Oxidative stress is a crucial driver of various pathomechanisms during OA progression, including catabolic and pro-inflammatory processes, regulated cell death, and senescence [14, 15, 18, 25]. To clarify whether ROS represent a potential trigger of GDF-15 expression, we investigated its expression after induction of oxidative stress in chondrocytes.

We previously reported about the harmful consequences of enhanced oxidative stress after cartilage trauma in a human *ex vivo* and rabbit *in vivo* model, which could be attenuated by antioxidative therapy using N-acetyl cysteine (NAC) [18, 19, 26]. In accordance with our assumption, addition of NAC significantly reduced the secretion of GDF-15 after *ex vivo* cartilage trauma as compared to the untreated impacted tissue (**Figure 4A**). Moreover, the gene expression and secretion of GDF-15 was significantly increased upon exposure of isolated hAC to H_2_O_2_ and consequent intracellular ROS accumulation (**Figure 4B-E**). In line with the results from the *ex vivo* cartilage trauma model, stress-induced secretion of GDF-15 was almost completely abolished by addition of NAC (**Figure 4E**). Taken together, these data imply that oxidative stress and subsequent cellular damage induces GDF-15 expression in hAC.

**Figure 4:**
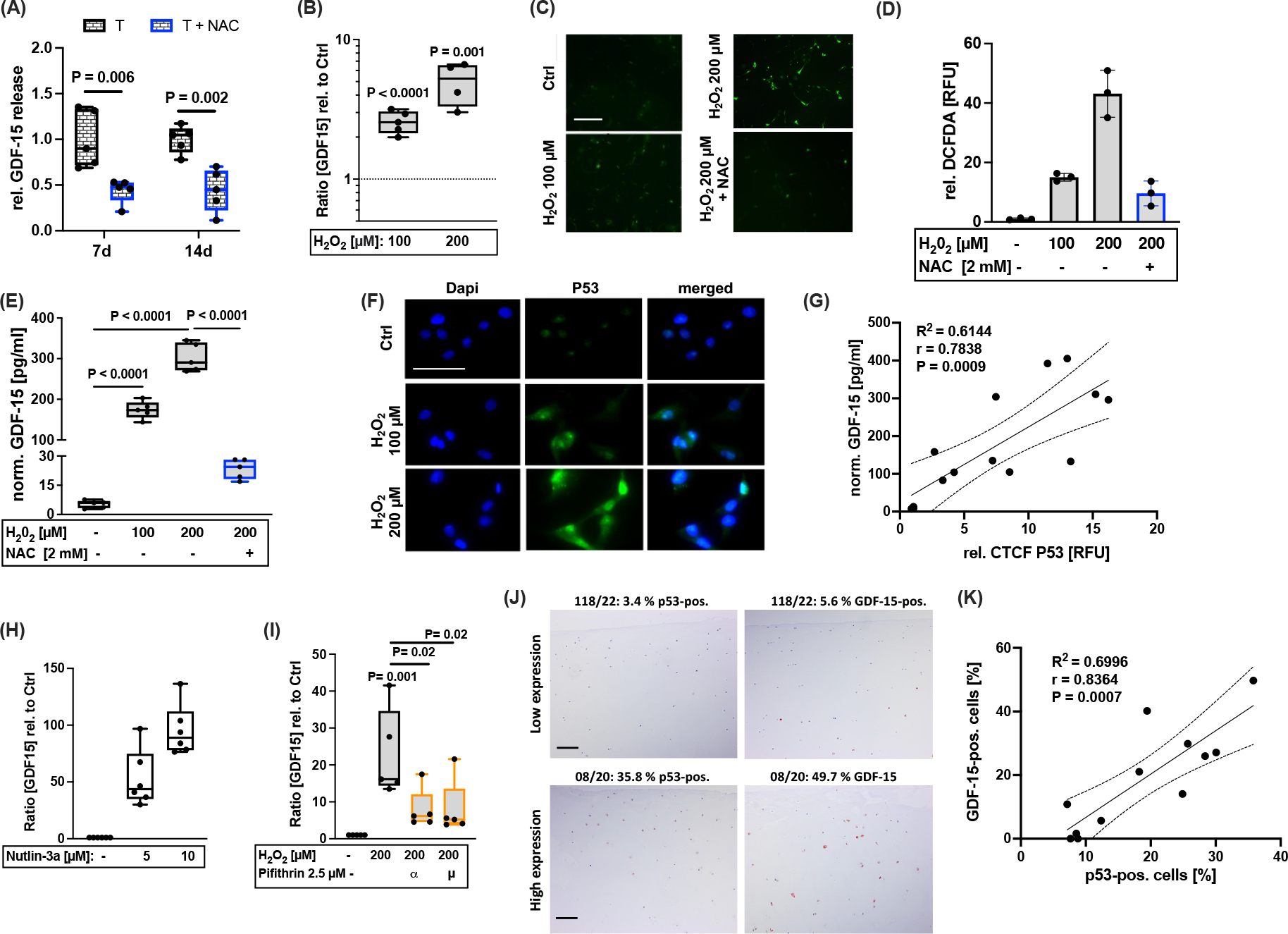
GDF-15 expression is modulated by ROS-induced p53 activation. (A) Relative release of GDF-15 at 7 d and 14 d after *ex vivo* cartilage trauma of untreated (T) or NAC-treated (T + NAC) explants. (B) Gene expression of GDF15 and (C,D) detection of intracellular ROS levels by DCFDA assay in isolated hAC upon H_2_O_2_ exposure for 48 h. (E) Release of GDF-15 by H_2_O_2_-treated hAC w/ or w/o treatment of NAC. (F) Representative images of IF staining of p53 in H_2_O_2_-treated hAC. (G) Correlation analysis between CTCF values of p53 staining and GDF-15 concentrations of unstimulated or H_2_O_2_-treated hAC. Gene expression analysis of GDF-15 in hAC after stimulation with (H) the MDM2 antagonist (p53 inducing) Nutlin-3a and (I) the inhibitor of p53 transcription activity, Pifithrin-α or Pifithrin-µ. (J,K) Representative images of IHC staining against p53 and GDF-15 in human cartilage and corresponding correlation analysis. The white bar in (C) and (F) represents 100 µm. The black bar in (J) represents 200 µm.

Oxidative stress has various effects regarding chondrocyte fate, comprising regulated cell death and senescence [18, 25]. Moreover, the tumor suppressor protein p53 is a key regulator of ROS-mediated cell fate decision and GDF-15 is considered as a transcriptional target of this stress-responsive transcription factor [14, 27]. To determine the role of p53 in GDF-15 expression in hAC, p53 activity was assessed by immunofluorescence (IF). As expected, p53 activity was significantly enhanced after exposure to H_2_O_2_ as indicated by enhanced fluorescence and nuclear translocation (**Figure 4F**). Subsequent correlation analysis of the p53 fluorescence signal (CTCF) and the release of GDF-15 in the corresponding samples revealed a positive association between the proteins (**Figure 4G**). Accordingly, induction of p53 activity by the MDM2 antagonist Nutlin-3a resulted in a concentration-dependent increase of the gene expression of GDF-15, while addition of the p53 inhibitors Pifithrin-α or Pifithrin-µ to H_2_O_2_-treated hAC significantly reduced the expression of the cytokine (**Figure 4H,I**). The positive correlation between p53 activity and GDF-15 expression was confirmed in human cartilage samples, comprising different OARSI grades as well as in impacted and unimpacted tissue sections (**Figure 4J-K**).

Taken together, GDF-15 expression in hAC is driven by oxidative stress and subsequent activation of the stress-inducible transcription factor p53.

### GDF-15 is secreted by senescent chondrocytes but does not induce paracrine senescence

Long-term exposure to non-cytotoxic oxidative stress and subsequent cell damage is considered as major driver of cellular senescence [14]. Recently, we described a novel doxorubicin (Doxo)-based *in vitro* model to reliably induce stable chondrosenescence in hAC [28]. We used this *in vitro* model to examine the expression of GDF-15 as a potential SASP factor of senescent chondrocytes. Chondrocyte senescence was confirmed by the expression of CDKN1A, CDKN2A, and p53 activity (**Figure 4A,B**). Moreover, senescent chondrocytes expressed high levels of GDF-15 (**Figure 4A,C**). To further confirm the association between chondrosenescence and GDF-15 expression, high passage OA chondrocytes were treated by the senolytics dasatinib and quercetin as previously described [29]. Selective elimination of senescent chondrocytes resulted in declined expression of *GDF15* and its transcription factor *p53* (**Figure 4D**).

As our findings indicated that hAC not only produce GDF-15 but might also act as recipient cells, in particularly under pathophysiologic conditions, we investigated the response of isolated hAC to exogenous rhGDF-15 by means of a bulk RNA sequencing. Comparison of rhGDF-15-stimulated hAC with untreated controls revealed that 93 genes were differentially expressed, with 40 genes being upregulated and 53 genes being downregulated (**Figure 4E, Supplement S3A**). By means of a Kyoto Encyclopedia of Genes and Genomes (KEGG) analysis, the regulated genes could be classified to cellular senescence (cellular processes) and glycerophospholipid metabolism among others (**Figure 4F**). Further analysis by Gene Ontology (GO) enrichment revealed that processes associated with biological regulation, metabolism, signalling, response to stimulus, development, and immune system were among the top regulated biological processes in hAC upon rhGDF-15 stimulation (**Supplement S3B-D**). However, we could not identify any significant effect of GDF-15 stimulation on the expression of genes related to SASP or cartilage matrix by RNASeq.

To further investigate the potential effects of secreted GDF-15 in terms of chondrosenescence, we performed additional qPCR analysis, focusing on senescence-associated target genes. Stimulation with rhGDF-15 for 48h reduced the gene expression of CDKN1A and CDKN2A, while having no effect on common SASP factors described in the context of OA [25, 28], such as CXCL1, IL-6, and MMP13 (**Figure S3E**). In line with the reduced mRNA levels of the cell cycle inhibitors, rhGDF-15 stimulation induced the proliferation of hAC, as determined by an alamarBlue assay (**Figure 5G**). Furthermore, addition of rhGDF-15 significantly increased the number of migrated cells in a scratch assay, which indicates elevated migratory and mitotic activity (**Figure 5H**). However, GDF-15 had no chemoattractive effect on mesenchymal stem cells, as investigated by means of a Boyden chamber assay (**Supplement S4A**). As proliferation and migration are commonly associated with tissue regeneration, we conducted an *in vitro* chondrogenesis assay on dedifferentiated hAC in presence of the cytokine. Addition of rhGDF-15 during chondrogenic re-differentiation of hAC decreased the overall score of neocartilage formation to some extent, but did not inhibit chondrogenesis per se (**Figure 5I,J**). Overall, these data imply that GDF-15 might serve as a SASP factor, which promotes migratory and mitotic properties in hAC, but does not accelerate chondrosenescence.

**Figure 5:**
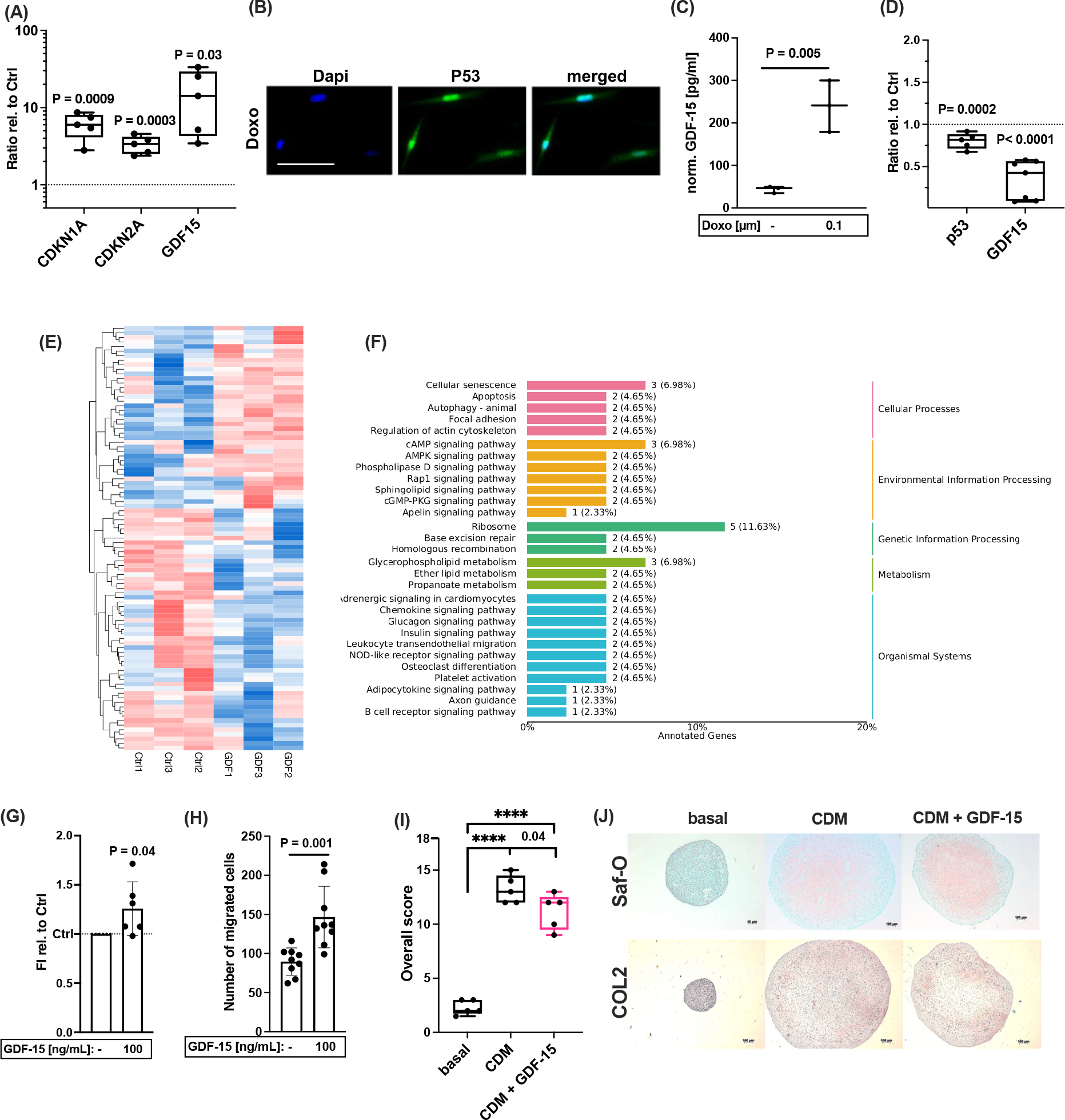
Response of hAC to exogenous GDF-15. Characterization of Doxo-stimulated hAC by (A) gene expression of CDKN1A, CDKN2A, and GDF15, and (B) IF of p53. (C) Release of GDF-15 by Doxo-stimulated hAC. (D) Gene expression of GDF15 and p53 at day 7 after selective elimination of senescent hAC by dasatinib (250 nM) and quercetin (5 µM) for 3d. (E) Heatmap presenting significantly upregulated (red) and downregulated (blue) genes after GDF-15 stimulation as compared to untreated control hAC. (F) GO enrichment analysis depicting classification of significantly regulated genes after GDF-15 stimulation as compared to untreated control hAC. (G) Relative fluorescence intensity (FI) of the alamarBlue assay 48h after stimulation of hAC with exogenous GDF-15. (H) Number of migrated hAC 48h after stimulation with GDF-15 in a scratch assay. (I) Scoring of in vitro redifferentiation of hAC after 4 weeks in chondrogenic differentiation medium (CDM) with or without addition of rhGDF-15. (J) Representative Safranin-O (Saf-O) and IHC staining against COL2 of respective redifferentiation. The white bar in (B) represents 100 µm. The black bar represents 100 µm.

### Exogenous GDF-15 mitigates trauma-induced pathomechanisms in human cartilage and might promote a pro-regenerative phenotype

To investigate the potential modulative effect of GDF-15 on hAC after cartilage injury, the cytokine was added for 7 d and 14 d after *ex vivo* trauma. While the mechanical impact reduced the cell viability as expected, addition of rhGDF-15 significantly prevented chondrocyte death at both time points (**Figure 6A,B**). Moreover, rhGDF-15 significantly reduced trauma-induced NO production after 7 d, whereas this effect could not be observed at the later time point (**Figure 6C,D)**. Although trauma-induced release of GAG was only increased by trend, treatment with rhGDF-15 significantly attenuated the amount of the matrix components in culture media of impacted cartilage explants (**Figure 6E,F**). Exemplary Safranin-O staining of human cartilage explants confirmed higher proteoglycan content in rhGDF-15-treated explants after *ex vivo* trauma and indicated enhanced cell cluster formation, regardless of a preceding impact (**Figure 6G**).

**Figure 6:**
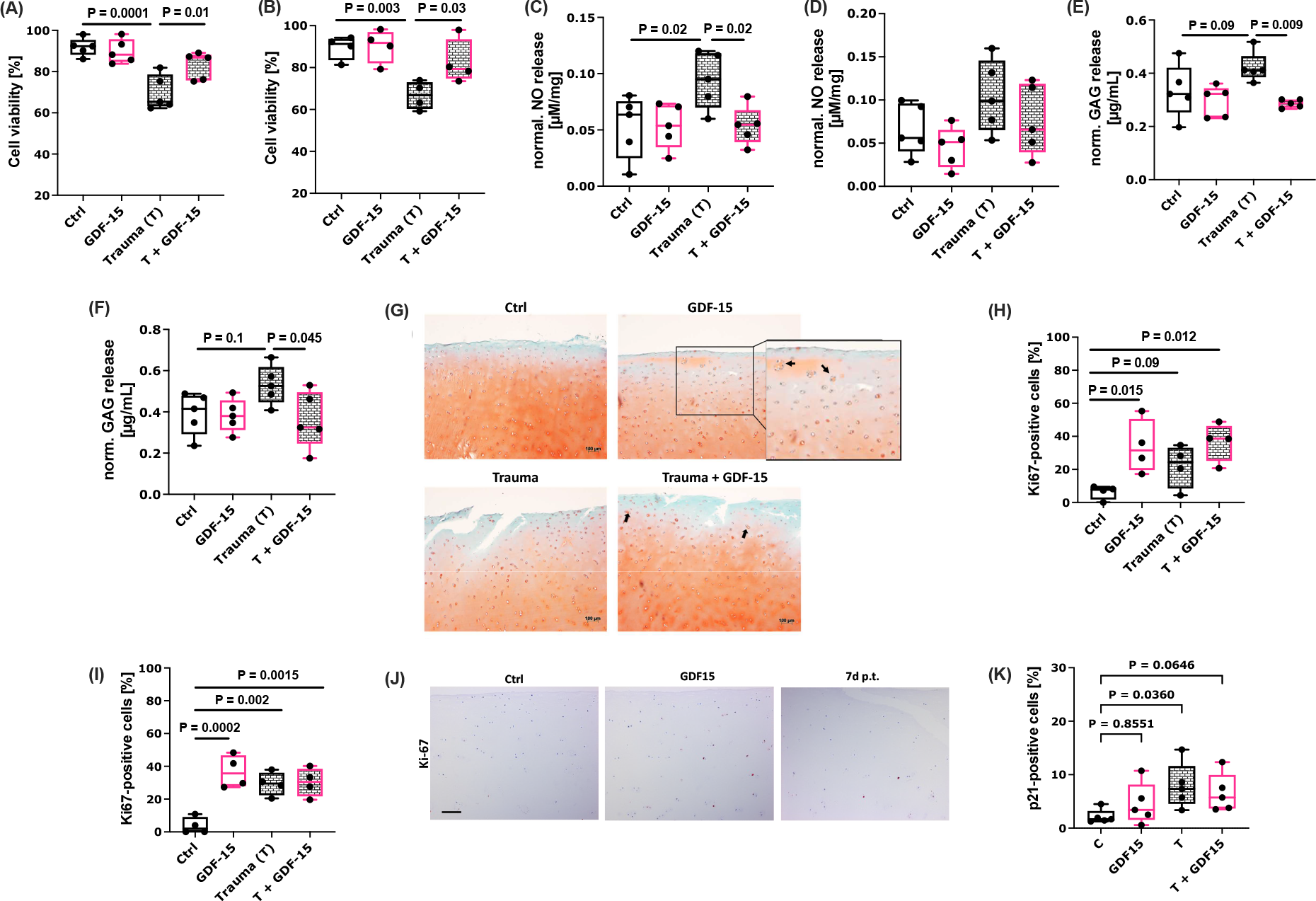
Effects of exogenous rhGDF-15 after ex vivo cartilage trauma. Cell viability of hAC at (A) 7 d and (B) 14 d after ex vivo cartilage trauma and/ or GDF-15 stimulation, determined by a live/dead assay. Normalized release of NO at (C) 7 d and (D) 14 d after *ex vivo* cartilage trauma and/ or GDF-15 stimulation, determined by a Griess assay. Normalized GAG release at (E) 7 d and (F) 14 d after *ex vivo* cartilage trauma and/ or GDF-15 stimulation, determined by a DMMB assay. (G) Representative Saf-O staining of human cartilage explants 7 d after *ex vivo* trauma and/ or GDF-15 stimulation. Percentage of Ki67-positive cells at (H) 7 d and (I) 14 d after *ex vivo* cartilage trauma and/ or GDF-15 stimulation. (J) Representative images of IHC staining against Ki-67 7 d after GDF-15 stimulation or *ex vivo* cartilage trauma. (K) Percentage of p21-positive cells at day 14 after *ex vivo* cartilage trauma and/ or GDF-15 stimulation. The black bar represents 200 µm.

To clarify potential pro-mitotic activity of chondrocytes upon rhGDF-15 stimulation, a Ki67 staining was performed. In line with the cluster formation described above, the percentage of Ki67-positive chondrocytes was significantly increased upon addition of rhGDF-15 in impacted and non-impacted cartilage explants. Interestingly, the pro-mitotic stimulus of exogenous rhGDF-15 was stronger than that of the cartilage trauma at day 7, whereas no difference could be found between traumatized explants w/ or w/o rhGDF-15 treatment after 14 d (**Figure 6H-J**). While we observed a significant increase of p21-positive chondrocytes at day 14 post trauma, long-term exposure to rhGDF-15 did not induce the expression of the cell cycle inhibitor in hAC (**Figure 6K,L**).

Moreover, we examined the immunomodulatory properties of rhGDF-15 in activated human macrophages (THP-1 cells). In accordance with the literature, rhGDF-15 mitigated the gene expression of TNF and IL1B to some extent but had no effect on IL6 (**Supplement S5A-C**). In hAC exposed to 10 ng/mL IL-1β, addition of rhGDF-15 restored the transcription of COL2A1 but had no effect on IL-1β-induced COX2 or MMP13 gene expression (**Figure S5D-F**).

Taken together, our results imply that exogenous rhGDF-15 has pro-regenerative and protective effects after cartilage injury without inducing chondrosenescence.

## Discussion

Traumatic injuries of joint-related tissues are considered to carry a high risk for development of a PTOA. However, the underlying pathomechanisms which are driving the pathogenesis are complex and not well understood yet.

In the present study we identified the stress-responsive cytokine GDF-15 as a potential indicator or even biomarker of PTOA in synovial fluid samples of end-stage OA patients. Further investigation revealed that GDF-15 expression was generally upregulated in OA cartilage tissue and highly induced after *ex vivo* cartilage trauma. Posttraumatic GDF-15 expression in chondrocytes was found to be triggered by oxidative stress and subsequent p53 activation. Although GDF-15 was associated with cellular stress and might represent a new SASP factor of senescent chondrocytes, the cytokine induced a pro-regenerative response, including cell proliferation, migration, and survival, in cartilage tissue.

Regarding our present findings, the strong correlation between syGDF-15 and cystatin C in PTOA and IOA patients might result from the fact that both GDF-15 and cystatin C are considered as common age- and disease-related biomarkers. Accordingly, we previously reported that sGDF-15 was significantly correlated with cystatin C and hs-CRP within the cohort of the Ulmer OA study, comprising 636 patients suffering from end-stage hip or knee OA [10]. Moreover, cystatin C and GDF-15 have lately been described among the three most significant plasma SASP markers in age and were associated with adverse health effects as demonstrated by various physiological parameters, including inflammation (IL-6 levels), gait speed, and grip strength [30].

The stress-responsive protein p53 is considered as a pivotal regulator in tissue regeneration as described in epithelial and myocardial repair as well as limb regeneration, among others [31-33]. In this context, p53 orchestrates cell fate decision and plasticity in a concentration-dependent manner. For one thing, low levels allow cell cycle re-entry of postmitotic cells and accumulation of progenitor cells, then again, high levels of p53 induce differentiation and transient senescence. In our study we observed enhanced GDF-15 expression upon ROS-induced p53 activation and subsequent senescence of chondrocytes. Usually, accumulation of senescent chondrocytes is considered as one of the major drivers of OA progression, due to the detrimental effects of the secreted SASP factors [14]. However, our findings demonstrated that addition of exogenous rhGDF-15 does not promote spreading of the senescent phenotype, but, on the contrary, induced proliferation, migration, and cell survival of hAC. Similar results were currently reported in case of IL-6, which is a common SASP factor and controversially discussed in cartilage degeneration and regeneration [34-36]. Overall, it is widely accepted that transient senescence contributes to tissue regeneration and that SASP factors initially accelerate related processes, such as immune and stem cell recruitment, fibroblast activation, and ECM remodeling [37]. In cartilage, transient senescence might become chronic due to the tissue-specific properties, in particularly hypocellularity and avascularity, resulting in a poor tissue regeneration and insufficient clearance of senescent chondrocytes. Consequent accumulation of senescent cells and continuing secretion of SASP mediators has emerged to be highly detrimental [13, 14]. Therefore, it cannot be completely ruled out that the pro-mitotic effect of GDF-15 in particularly in combination of other pro-inflammatory SASP factors and oxidative stress may switch from a pro-regenerative to an adverse effect, as previously demonstrated for pro-mitotic TGF-ß1 and FGF2 in combination with irradiation-induced DNA damage [38].

Besides the fact, that cellular senescence is not per se a pathophysiologic process, it should be noted that GDF-15 has been described as an anti-inflammatory and immunomodulatory cytokine. In line with this, a previous case-control study, including 910 RA patients and healthy controls, identified two GDF-15 gene polymorphisms, which were associated with an enhanced risk for RA [39]. They hypothesized that the immunomodulatory nature of GDF-15 might possess anti-arthritic effects. In fact, the cytokine was assumed to attenuate CXCL10/CXCR3-dependent infiltration of T lymphocytes during glomerulonephritis [40]. Accordingly, GDF-15 was found to impair dendritic cell maturation and thus their ability to serve as antigen-presenting cells during T-cell priming [41]. Its role as immunomodulatory cytokine might also explain that serum levels of GDF-15 were reported to correlate with disease activity in RA patients [42]. In our study, we could confirm a positive association between the inflammation marker hs-CRP and syGDF-15 levels in the PTOA group, an increased secretion of GDF-15 by TCM-stimulated hFLS, and anti-inflammatory effects of rhGDF-15 on activated THP1 to some extent.

Strikingly, we observed that hAC respond to exogenous rhGDF-15 and thus provided first evidence of GFRAL, the only member of the GNDF family receptor-α (GFRα) capable of binding GDF-15 [43], in cartilage tissue. This implies that GDF-15-mediated effects result from its interaction with GFRAL. As we found enhanced levels of GFRAL in highly degenerated and traumatized cartilage tissue, we assume an increased susceptibility of chondrocytes to the cytokine in the course of disease progression and trauma response. Although it has been initially hypothesized that GFRAL expression was restricted to the central nervous system [44], more recent studies report the presence of GFRAL in peripheral tissues, such as adipose, pancreatic ductal adenocarcinoma, and gastric cancer tissue [45-47]. Moreover, we excluded that the response of hAC towards exogenous rhGDF-15 was mediated via interaction with the TGF receptor or ErbB2, which both have been discussed as alternative receptors of GDF-15 [48, 49], by addition of the corresponding inhibitors in migration assays (**Supplement S4B**,**C**).

With regards to the current knowledge about GDF-15, which is also described as a moonlight protein due to its various, independent functions beyond originally identified ones [50], we are convinced that our results display only a small range of the comprehensive role of GDF-15 in cartilage biology and pathophysiology. Although our findings are based on human material, thus providing valuable information about the clinical situation, further *in vivo* studies might be needed to understand the role of GDF-15 in age-related and posttraumatic OA development. Evidence of potentially detrimental effects of GDF-15 could not be found in this study, but cannot be fully excluded as mentioned above. Moreover, the potential suitability of GDF-15 as a prognostic or diagnostic biomarker in PTOA has to be further investigated in a long-term study on patients suffering from joint injuries.

Altogether, the current study provides first evidence for the p53-mediated expression of GDF-15 in chondrocytes upon cellular stress and its potential cell and chondroprotective effects after *ex vivo* cartilage trauma. Although GDF-15 expression was induced by the transcription factor p53 and was linked to cellular senescence, we could not observe any adverse impact of the cytokine on cartilage homeostasis. Regarding our findings, we conclude that GDF-15 might represent a novel marker of PTOA and might initially promote pro-regenerative processes. Whether GDF-15 may serve as a future target in OA therapy has to be clarified in subsequent studies.

## Author contribution

JR: funding acquisition, study concept and design, data acquisition, interpretation of data, figure preparation, statistical analysis, and writing/editing of the manuscript. SSM: data acquisition, conducting experiments, and participation in manuscript preparation. AL, SM: data acquisition, conducting experiments, and revision of manuscript. RB, DR: administration Ulm OA study, data interpretation, and revision of the manuscript. All authors approved the final version of the manuscript and ensure that questions related to the accuracy or integrity of any part of the work are appropriately investigated and resolved.

## Acknowledgments

The authors would like to thank Natalie Braun and Christiane Schulz for excellent technical assistance. This research study was supported by the European Social Fund and by the Ministry of Science, Research and Arts Baden-Württemberg as well as the University of Ulm (Hertha-Nathorff-Programm).

