## Supplement for "p53-induced GDF-15 expression promotes a pro-regenerative response in human chondrocytes upon cartilage injury"

#### Table S1: List of primer and TaqMan® gene expression assays used for qRT-PCR.

| Target gene | MasterMix | Primer sequence/ TaqMan Probe |
|---|---|---|
| 18S rRNA | Power SYBR® Green PCR Master Mix (Applied Biosystems) | 5′-CGCAGCTAGGAATAATGGAAT AGG-3’ (forward) and 5′-CATGGC CTCAGTTCCGAAA-3’ (reverse) |
| CDKN1A (P21) | TaqMan® fast advanced MasterMix | Hs00355782 |
| CDKN2A (P16INK4/P14ARF) | | Hs00923894 |
| COL2A1 | | Hs00264051 |
| CXCL1 | | Hs00605382 |
| ERBB2 | | Hs01001580_m1 |
| GAPDH | | Hs02758991 |
| GAPDH | Platinum® SYBR® Green qPCR SuperMix-UDG (Invitrogen, Darmstadt, Germany) | 5′-TGGTATCGTGGAAGGACTCATG-3’ (forward) and 5′-TCTTCTGGGTGG CAGTGATG-3’ (reverse) |
| GDF15 | TaqMan® fast advanced MasterMix | Hs00171132_m1 |
| GFRAL | | Hs01087628_m1 |
| HPRT1 | | Hs02800695 |
| IL1B | | Hs00174097_m1 |
| IL6 | | Hs00985639 |
| MMP-13 | | Hs00233992 |
| TMEM199 | | Hs01022209 |
| TNF | | Hs01113624_g1 |
| TP53 | | Hs01034249_m1 |

#### Supplement S1: Influence of privious hyaluronidase digestion on concentrations of synovial GDF-15.

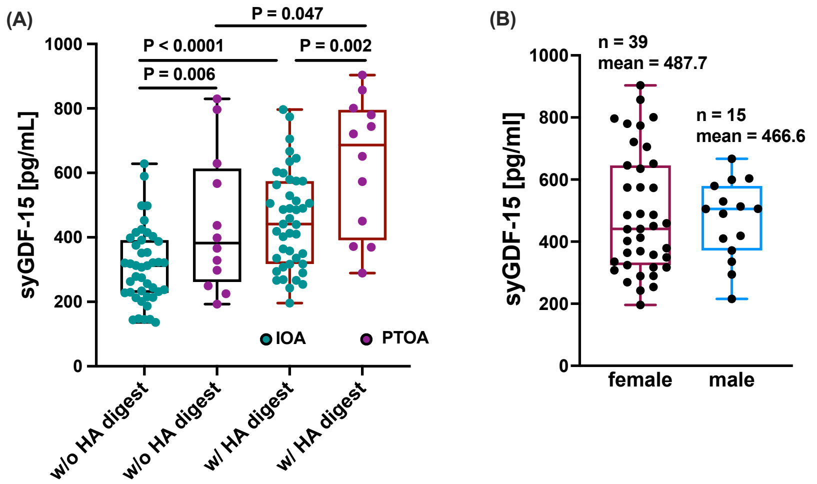


(A) Comparison between syGDF-15 quantification with (w/) or without (w/o) preceding hyaluronidase digestion. (B) Comparison of syGDF-15 concentrations in female and male OA patients (IOA and PTOA combined).

**Supplement S2: Results of the gene expression analysis of GDF-15 and GFRAL after cartilage trauma.**

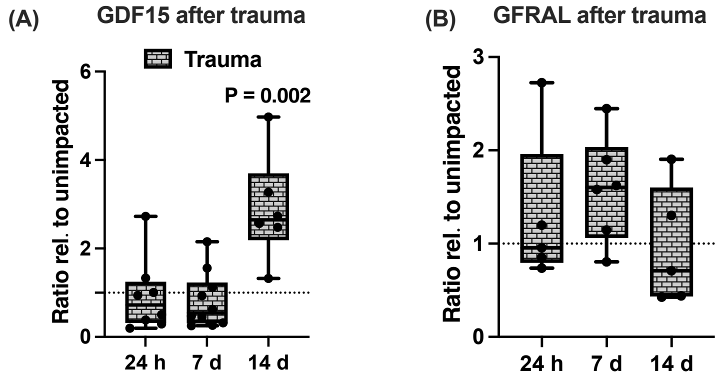


Gene expression analysis of (A) GDF15 and (B) GFRAL at 24 h, 7 d, and 14 d after *ex vivo* cartilage trauma.

**Supplement S3: Additional findings of RNA-seq analysis and confirmation of senescence-associated genes by means of qRT-PCR.**

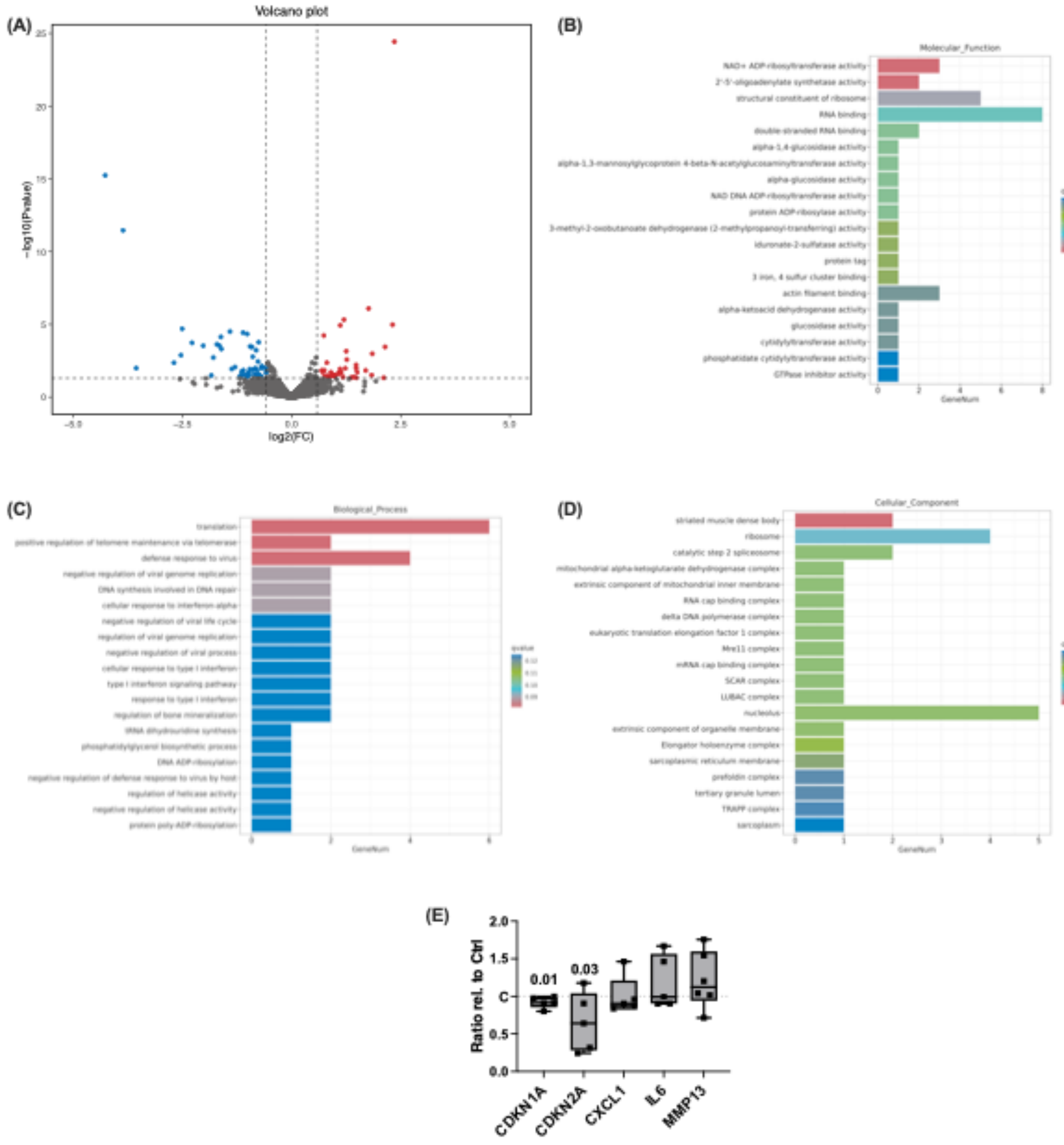

(A) Volcano blot of differentially expressed genes in hAC after GDF-15 stimulation. Gene Ontology (GO) enrichment of significantly regulated genes in response to rhGDF-15; top terms classified as (B) molecular function, (C) biological process, and (D) cellular components. (E) Gene expression analysis of senescence-associated markers 48 h after GDF-15 stimulation.

### Supplement S4: Influence of GDF-15 on directed (boyden chamber) and non-directed (scratch assay) migration

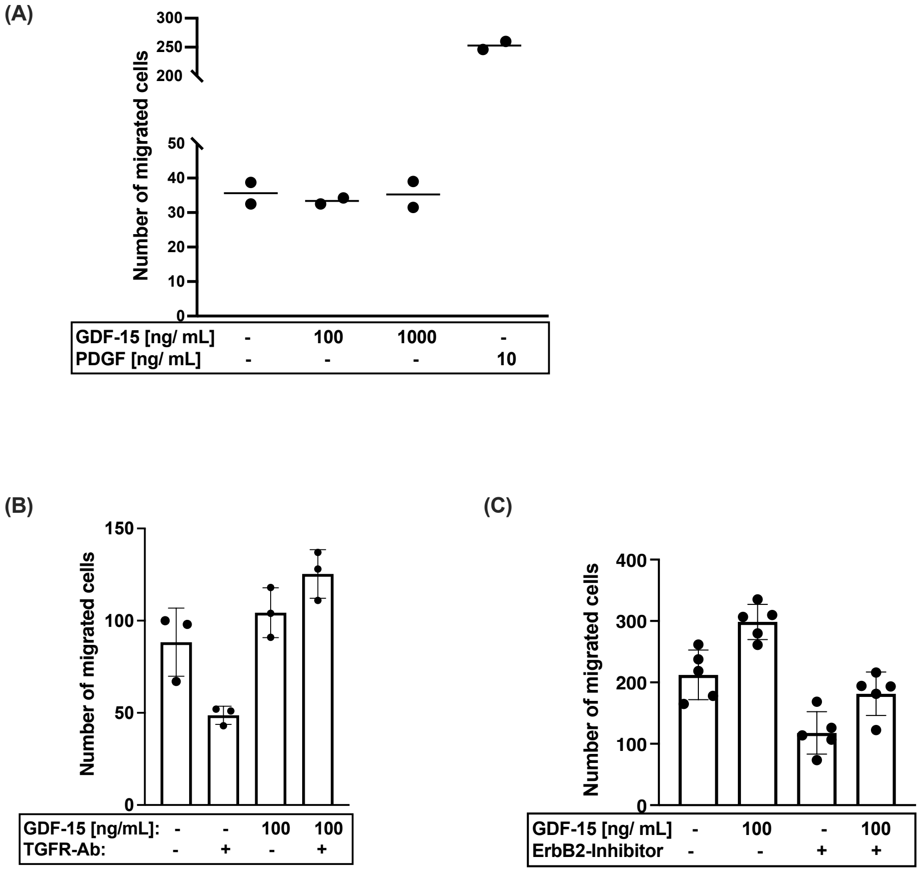


(A) Boyden chamber assay of human mesenchymal stem cells in response to rhGDF-15 or PDGF as positive control (PK). Scratch assay of hAC in response to (B) rhGDF-15 w/ or w/o inhibitory TGF receptor antibody or (C) ErbB2 inhibitor.

### Supplement S5: Anti-inflammatory/ immunomodulatory effects of rhGDF-15.

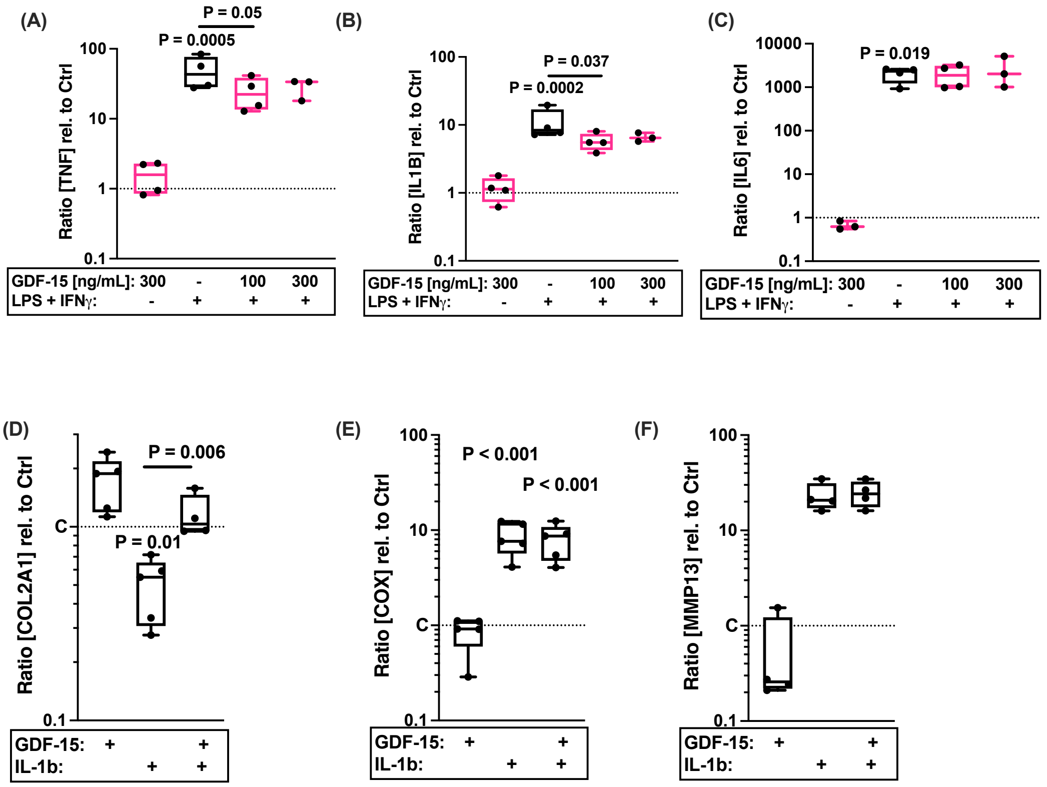


Gene expression of (A) TNF, (B) IL1B, and (C) IL6 in 100 ng/ mL LPS + 20 nM IFNy-stimulated THP-1 cells w/ or w/o addition of GDF-15. Gene expression of (D) COL2A1, (E) COX2, and (F) MMP13 in IL-1β-stimulated hAC w/ or w/o addition of GDF-15.
